# Gut microbiome diversity associates with estimated lifetime and annual reproductive success in male but not female collared flycatchers

**DOI:** 10.1101/2024.08.12.607561

**Authors:** Liukkonen Martta, Gustafsson Lars, Grond Kirsten, Ruuskanen Suvi

## Abstract

The gut microbiome (hereafter, GM) varies across individuals of the same species and this pattern has been observed in multiple wild species. Evidence shows that the GM connects to individual health and survival especially in captive species, but more research is needed to understand how the GM connects to host fitness in wild species. We used long-term monitoring data to investigate whether the GM of collared flycatchers *Ficedula albicollis* associates with annual and lifetime reproductive success (LRS), and survival to the following breeding season. This is the first study that 1) characterized the collared flycatcher GM, and 2) investigated how variation in the GM related to LRS in wild birds. Our results showed that higher GM diversity was associated with a higher annual and lifetime reproductive success in especially male collared flycatchers. We also found that the compositional variation in collared flycatcher GMs was explained by sex, age, and breeding habitat, but not by annual or lifetime reproductive success. Individuals that died before the next breeding season had higher abundances of ASVs belonging to the pathogenic families *Enterobacteriaceae* and *Parachlamydiaceae*, and the genera *Corynebacteria* and *Sphingomonas*. Our results show that the GM associates with different aspects of host fitness in a wild bird population. More research is needed to evaluate if there is a causal relationship between the GM and individual fitness. These findings also contribute to our understanding of the GMs role in evolution by elucidating the connection between the GM (trait) and reproductive success.

## 1 INTRODUCTION

Lifetime reproductive success (hereafter, LRS) is a commonly used measure of individual fitness in natural populations, and it measures the total number of offspring that an individual produces during their life (Arnold and Wade, 1984, Clutton-Brock et al., 1988, Newton 1989). Survival to following breeding season and annual reproductive success (hereafter, ARS) are the most important determinants of LRS, because they define the number of offspring an individual produces during their life (Murray, 2000). Factors such as behavior (Buzatto et al., 2007; Réale et al., 2009), parental morphological traits (Fox et al., 1995; Grant and Grant, 2000; Simmons, 1988), and the time of breeding, hatch date or parental condition (Part and Gustafsson, 1989a; Verhulst et al., 1995) can influence LRS. For example, in wild European rabbits *Oryctolagus cuniculus*, social rank largely influenced LRS; In high-ranking female rabbits LRS and survival were approximately 60 % greater than in lower-ranking females (von Holst et al., 2002). In great tit *Parus major* and house sparrow *Passer domesticus*, adult individuals with higher body mass and longer wing and tarsus length produced higher quality offspring that were more likely to survive than their lower body condition counterparts (Jensen et al., 2004; Pigeault et al., 2020). Similar results have been observed in the collared flycatchers (Gustafsson, 1989).

Birds make an excellent study system for investigating the factors that influence LRS as long-term individual monitoring is possible in many species. Multiple studies have investigated LRS and the various factors contributing to LRS such as fledging date (Visser and Verboven, 1999), ornament display (Costanzo et al., 2017; Potti et al., 2013), parental condition and age (Harvey et al., 1985; Schroeder et al., 2015), male quality (Costanzo et al., 2017), and telomere length (Sudyka et al., 2019). One factor that has not been studied in wild bird populations but may associate with LRS and influence survival and ARS, is the gut microbiome (GM) (as reviewed in Comizzoli et al., 2021).

The vertebrate GM consists of a complex community of bacteria, archaea, viruses, microbial eukaryotes, and their genomes (Ley et al., 2008). It is a vital part of host biology and health due to its importance in growth, nutrient absorption, metabolism, immune system functioning (Amato, 2016; Kau et al., 2011; McFall-Ngai et al., 2013), pathogen susceptibility (Worsley et al., 2021), and behavior (Davidson et al., 2020). The GM is also a key part of the gut-brain axis i.e., the interaction between individual’s gut and brain, and therefore can also affect individual stress responses and behavior (Chen et al., 2013; Cryan and O’Mahony, 2011). Generally, high GM diversity is usually more stable and may indicate better host health (Lozupone et al., 2012). However, the GM can fluctuate during the host’s lifespan and can rapidly change in response to environmental factors (Candela et al., 2012). Some of these fluctuations result from the GM adapting to environmental change and some studies have shown how the GM can be shaped by factors such as diet (Baniel et al., 2021; Davidson et al., 2020; Gong et al., 2021; Góngora et al., 2021), habitat (Drobniak et al., 2022; Loo et al., 2019), and season (Davenport et al., 2014; Escallón et al., 2019).

Because of its key role in supporting individual physiological functions such as digestion, immune system functioning, and metabolism throughout an individual’s lifespan, the GM has been suggested to affect individual survival (Rosshart et al., 2017; Sharpton, 2018). In the water flea *Daphnia magna*, a reciprocal GM transplant was shown to improve host stress tolerance and mediate local adaptation, suggesting that GM composition can influence survival (Houwenhuyse et al., 2021). In the genetically modified diabetic *db/db* mice intermittent fasting changed the GM composition leading to a higher survival rate compared to *ad lib* fed individuals (Beli et al., 2018). So far, the first (and only) study investigating association between the GM and fitness in a wild bird population showed that in adult Seychelles warblers *Acrocephalus sechellensis* individuals that did not survive into the following breeding season had an increased abundance of known pathogenic bacteria in their gut (Worsley et al., 2021). Studies investigating the GM’s link to survival in wild nestling birds are limited and the results are mixed. The GM of nestling great tits did not associate with survival to fledging (Liukkonen et al., 2023), whereas there was a time-lagged association between GM diversity and nestling condition: higher abundance of specific *Lactobacillus* taxa abundance positively correlated with nestling weight and survival (Davidson et al., 2021).

Besides survival, the GM is likely to influence the reproductive success component of LRS, both directly by influencing reproductive physiology, and indirectly by influencing parental body condition. Firstly, the GM regulates hormone function and stimulates reproductive hormones, such as estrogen, progesterone, and testosterone (as reviewed by Hussain et al., 2021). Secondly, reproduction is energetically costly (Nilsson and Svensson, 1997). Low body condition, which may be a result of disruptions in the GM, can influence reproductive success (Ben-Yosef et al., 2008; Morimoto et al., 2017). Indeed, a few studies in captive and wild animals have reported associations between the GM and reproduction. In eastern black rhino *Diceros bicornis michaeli* the GM composition was significantly associated with breeding success and approximately a third of the observed bacterial genera correlated with reproductive hormone concentrations (Antwis et al., 2019). In fruit flies *Drosophila melanogaster,* a reciprocal GM transplant changed mating duration in males, increased short-term offspring production in females, and contributed to the female offspring’s GM (Morimoto et al., 2017). In one of the few bird studies investigating the hormone-GM relationship, yellow-legged gull *Larus michahellis* glucocorticoid levels were shown to influence the GM composition. Corticosterone-implanted male and female birds had reductions in both pathogenic and potentially beneficial taxa implying links between the GM and avian neuroendocrine system (Noguera et al., 2018). In the endangered crested ibis *Nipponia nippon*, there was an association between both GM diversity and composition and reproductive status between sterile and healthy ibises, which could reflect on ARS and LRS as well (Ran et al., 2021). All in all, there could be a (sex-specific) link between the GM and reproductive success in birds as well.

In this study, we use long-term monitoring data to explore whether the bacterial GM of wild adult collared flycatcher males and females associates with their survival and reproductive success and ultimately LRS. We analyzed whether there is a correlation between 1) lifetime reproductive success (LRS) and the gut microbiome, 2) annual reproductive success (ARS) and the gut microbiome, and 3) survival to the following breeding season and the gut microbiome. We predict that higher GM diversity positively correlates with both LRS and ARS, because high GM diversity is usually more stable and indicates better host health (as reviewed in Lozupone et al., 2012), and could possibly reflect to reproductive success as well. Furthermore, we predict that there are compositional differences in beneficial and pathogenic bacterial taxa in the GM of individuals that survive to the following breeding season vs. those that do not survive. To our knowledge, this is the first study on the association between lifetime reproductive success and the GM in a wild population of birds. We expect this study to be of interest to a wider scientific audience as it sheds a light on the GMs link to reproduction, which is at the core of evolution.

## 2 METHODS

### 2.1 Study species

Collared flycatchers are small (∼ 13 g) migratory passerine birds that nest in cavities and will use artificial nest boxes if provided. The birds arrive from their wintering areas in southern Africa to the breeding areas in late April to mid-May. The collared flycatchers on Gotland (Sweden) lay one clutch per year, consisting of six eggs on average (range 4-8). At the beginning of May, the first eggs are laid, and incubation begins, a task which is undertaken solely by females. Collared flycatchers are socially monogamous birds, but extra pair copulations (EPC) and extra pair paternity (EPP) are frequently observed in the species. Nestlings hatch after approximately 14 days of incubation, remain in the nests for an additional 14-16 days, and are fed by both parents. They reach the maximum body mass at the age of 10-11 days and lose some mass before fledging. Then, fledglings stay close to the nest for another two weeks and are still fed by their parents. Monitoring of the population biology of the collared flycatcher has been carried out at Gotland, Sweden (57°03′ N, 18°17′ E) for 44 years (1980–2023). The birds breed in deciduous forests and prefer nest boxes over natural tree holes (Gustafsson, 1986), which makes them easy to handle. Collared flycatchers present an ideal study species for addressing our objectives due to their high site fidelity, the advantage nest box use provides for trapping and nestling monitoring. The collared flycatchers breeding at our study site exhibit high site fidelity and return rates of adults and young, (Gustafsson, 1986; Part and Gustafsson, 1989b) enabling us to study LRS, ARS, and survival to following breeding season.

### 2.2 General procedures to collect data on reproduction and survival

In the studied nest box population of the collared flycatcher, birds were checked weekly to gather breeding data including laying date, clutch size, and the number of fledglings and recruits. Females were trapped at the nest during incubation and males were caught while feeding nestlings (May-June). After catching, each bird was banded with a metal band, aged 1 year or older (Svensson 1992). All breeding attempts were carefully monitored until the chicks fledged. On day 12 after hatching, nestlings were measured, weighed, and banded with a metal band. In the study, we used the longitudinal data set, containing all the records on annual reproductive performance (i.e., the number of fledglings) of each bird used in this study and used this as an estimation of LRS.

### 2.3 Gut microbiome sample collection

All fecal samples were collected from wild adult females and males during breeding season in the summer of 2015. For LRS we use this one-year fecal sample data as a proxy of lifetime differences in GM across individuals. Current knowledge suggest that the GM is relatively stable during adulthood in wild birds and is quite robust to variation in the environment (Kreisinger et al., 2017; Somers et al., 2023). We collected fecal samples by placing each bird into a lined paper bag for 5-10 minutes until defecation (Knutie and Gotanda, 2018). Samples were stored on ice onsite and then moved to -80°C freezers for long-term storage. Each bird was aged, weighed, and had their tarsus and wing length measured. Body condition of the birds was based on the residuals of body mass on tarsus length (as done in e.g., Hemborg and Lundberg, 1998; Potti, 1999; Rosivall et al., 2009). Birds were aged based on the original ringing data as most birds were first ringed on Gotland Island as nestlings. The age of each bird was estimated based on these records and birds were categorized into three age groups: 1-year-old (age group 1), 2-4-year-olds (age group 2), and 4+ years old (age group 3). When unsure whether the bird was e.g., 1 or 2 years old we marked it as 1.5 years old. Furthermore, we used monitoring data from years before and after 2015 to estimate each flycatcher’s LRS, ARS and survival to following breeding season post-2015. We defined LRS based on the sum of all successfully fledged nestlings in all the years the adult bird was caught breeding on the island of Gotland (as done in Bouwhuis et al., 2015; Wysocki et al., 2019). ARS was the number of successfully fledged nestlings in 2015.

### 2.4 DNA extraction to sequence the bacterial 16S rRNA gene

DNA was extracted from fecal samples with Qiagen’s QIAamp PowerFecal Pro DNA Kit (Qiagen, Hilden, Germany). We used the manufacturer’s protocol with the following adjustments: a 10-minute incubation at 65 °C prior to lysis step was added and DNA eluent was filtered twice at the end to maximize DNA yield. Samples were randomized across extractions. A negative extraction blank was added to each DNA extraction batch to control for contamination during extraction. Post-extraction, the V4 region of the 16S rRNA gene (∼254 bp) was amplified using the primers 515F (5’-GTGYCAGCMGCCGCGGTAA - 3’) (Parada et al., 2016) and 806R (5’-GGACTACNVGGGTWTCTAAT – 3’) (Apprill et al., 2015). PCR reaction volume was 12 microliters and MyTaq RedMix DNA polymerase (Meridian Bioscience; Cincinnati, OH, USA) was used in the reactions. PCR protocol was 1) an initial denaturation at 95 °C for 3 minutes 2) 30 cycles of 95 °C for 45 sec., 55 °C for 60 sec., and 72 °C for 90 sec., and 3) a 10-minute extension at 72 °C at the end. To attach the Illumina barcodes for sample identification, a second round of PCR was done with 1) initial denaturation at 95 °C for 3 minutes, 2) 18 cycles of 98 °C for 20 sec., 60 °C for 15 sec., and 72 °C for 30 sec., and 3) final extension at 72 °C for 3 minutes. Each PCR plate had a negative control (nuclease free H_2_O) and a ZymoBIOMICS community standard (Zymo Research Corp, Irvine, CA, USA) to control for contamination during PCR and to confirm successful amplification. DNA concentration of the PCR products was measured with Quant-IT PicoGreen dsDNA Assay Kit (ThermoFischer Scientific, Waltham, MA, USA) and quality was inspected with gel electrophoresis (1.5 % TAE agarose gel). PCR products were equimolarly pooled and purified using the NucleoMag NGS Clean-up and Size Select beads (Macherey-Nagel; Düren, Germany). The pools were sequenced with Illumina Novaseq 6000 x 250 bp (San Diego, CA, USA) at the Finnish Functional Genomics Center and the University of Turku (Turku, Finland).

### 2.5 Sequence processing

The demultiplexed sequences were processed with QIIME2 (Bolyen et al., 2018) following the developer’s instructions for the 16S rRNA gene V4 region protocol. First, the Illumina adapters were removed with Cutadapt plugin version 4.4 (Martin, 2011) and sequence quality scores were inspected. DADA2 plugin version 2021.4.0 (Callahan et al., 2016) was used to truncate reads at 220 bp and to generate amplicon sequence variants (ASVs) that refer to each unique identified sequence (Eren et al., 2013). Taxonomy was classified by training a naïve-Bayes classifier on the SILVA v132 reference (Quast et al., 2013; Yilmaz et al., 2014). The phylogeny plugin was used to construct a rooted phylogenetic tree (Katoh et al., 2002). Eukaryotes, mitochondria, archaea, chloroplasts, and unassigned taxa were removed in QIIME2 as they were not of interest in this study. Sequences present in the DNA control samples and PCR negative controls were removed from the sample data to remove contaminants with the R package *decontam* (version 1.12; Davis et al., 2018). Also, singleton reads were removed as they are likely to be contaminant sequences or have minimal influence in this study. A total of 894 ASVs were filtered out as possible contaminants. Furthermore, ASVs with fewer than 100 reads across all samples were filtered out because of likely contamination. A total of 22 632 ASVs across 187 samples remained (SI 2). Resulting ASV data was then combined with the metadata, taxonomy table and phylogenetic tree with the *phyloseq* (McMurdie and Holmes, 2013) package version 1.44.0 in the R program (v 4.3.0, R Core Team).

### 2.6 Statistical Analyses

#### 2.6.1 Gut microbiome (alpha) diversity

To control for variation in sequence library size across samples, samples were rarefied to a depth of 2000 reads (Cameron et al., 2021; McMurdie and Holmes, 2014). Rarefying resulted in removal of two samples from the dataset, leaving 185 samples. We use Shannon Diversity Index (the abundance and evenness of taxa) and Chao1 Richness (estimation of bacterial richness per sample) as to measure GM alpha diversity. Shannon Diversity Index (hereafter, Shannon) is a commonly used metric to measure diversity. It is not affected by the presence of rare taxa like Chao1 (hereafter, Chao1) is and thus, making it more robust (Haegeman et al., 2013).

To test whether GM diversity in the year of sample collection associated with LRS and ARS, we used linear mixed effects models from the lme4 package (version 1.1-34, Bates et al. 2014). We used LRS and ARS as response variables, and Shannon/Chao1, body condition, age, and hatch date as explanatory variables. Study plot was included as a random factor. We first ran each model with all birds and then each sex i.e., females and males, separately. We did this because we had an unequal number of sexes (N_females_=122, N_males_=63) in our data and because there are sex specific differences in collared flycatcher LRS and survival (Herényi et al., 2014; Part et al., 1992), and these could reflect to the GM as well. We included body condition and age group in each statistical model because both can potentially influence LRS and ARS (Cichoñ, 2003; Cichoñ et al., 1998; Sendecka et al., 2007). Birds were categorized into age groups (3 groups: 1 year, 1.5-4 year, and 4+ year) because first time breeders (1-year-olds) are inexperienced in tending to their young and this may affect their reproductive success. Middle-aged birds (1.5 to 4-year-olds) were assigned as one group and old birds (4+ years old) in one group (Cichoñ, 2003; Sendecka et al., 2007). We also included hatch date as a proxy for the time of breeding in the models in which LRS or ARS are used as the response variable because hatch date is a known contributor to both LRS and ARS (Verhulst et al., 1995). Breeding area is included as a random effect in each model to control for sampling in the same area. In the models with LRS, we excluded samples (N_females_=26, N_males_=17) that were from the area PA because LRS was not monitored in those areas post-2015. To test whether GM diversity associated with survival to following breeding season, we used generalized linear mixed models from the *lme4* package (version 1.1-34, Bates et al. 2014). Survival (yes-no) was used as the response variable, and Shannon/Chao, body condition index, and age were included as explanatory variables. Study plot was included in the model as a random factor to control for samples coming from same nest box areas, which differ somewhat e.g., in habitat type. Same as with the models in which LRS is the response variable, we excluded samples (N_females_=26, N_males_=17) that were from the area PA because LRS was not monitored in those areas post-2015.

To further understand how GM diversity may be further influenced by environmental and intrinsic factors (that may be linked with LRS/ARS or survival), we ran an additional linear mixed effects model in which we analyzed whether sex, body condition, age group and plot area contributed (random) to GM diversity (response). We ran the model first with both sexes and then separately for each sex.

The significance of the factors in each model was tested using F-test ratios in analysis of variance (ANOVA). To check for multicollinearity between factors, we calculated variance inflation factors (VIFs<4) for each model with the *DHARMa* package (version 0.4.6 Hartig and Hartig, 2017). Results were plotted using *ggplot2* version 3.4.4 (Wickham et al., 2016).

#### 2.6.2 Gut microbiome composition (beta diversity)

We used the unrarefied dataset in GM composition (i.e., beta diversity) analyses and excluded the samples that had lower than 2000 reads to match the sample dataset with the GM diversity analyses. We calculated composition metrics using the *phyloseq* package (version 1.41.1, McMurdie and Holmes, 2013) and visualized composition with non-metric multidimensional scaling (NMDS) using Bray-Curtis dissimilarity metric, which is robust to homogeneity assumption violations (Anderson and Walsh, 2013; Schroeder and Jenkins, 2018). For plotting we used ggplot2 version 3.4.4 (Wickham et al., 2016). To assess the differences in GM composition between groups of samples, we ran a permutational multivariate analysis of variance (PERMANOVA) with 9999 permutations using the *adonis2* function from the package *vegan (*version 2.6.-4, Oksanen et al. 2008). We used the *betadisper* function to check for the homogeneity of variances in group dispersion.

First, to identify compositional differences in the GM of individuals of different LRS / ARS, we conducted a PERMANOVA with 9999 permutations. LRS / ARS was set as a predicting variable and sex, body condition, age group, breeding area, and hatch date were set as additional predictors. Second, to investigate compositional differences in GM between individuals that survived and individuals that did not survive to the following breeding season, we ran a PERMANOVA with 9999 permutations. Survival to the following breeding season (yes-no), sex, body condition, age group, and breeding area were included as predictors. To account for the size difference in the number of samples between sexes, we ran two additional models with only female and male samples. In these sex-specific models, survival to the following breeding season, age group, area, and hatch date were set as predictors contributing to GM composition.

#### 2.6.3 Differential taxa abundance

To investigate differential abundances in microbial taxa of the individuals that either survived or did not survive to the following breeding season, we ran a differential abundance analysis (DESeq2) with the DESEq2 package for the whole dataset and for the sexes separately (version 1.40,1; Love et al., 2016). A cut-off of P_adj_<0.05 was used to assess significance. Differentially abundant taxa were identified to the lowest possible taxonomic depth and visualized at Family level to improve readability.

## 3 RESULTS

### 3.1 Collared flycatcher gut microbiome

The unrarefied dataset had a total of 33,726,887 reads (range per sample: 260-853,017; average per sample: 170,338) that were assigned to 22,632 individual ASVs. There were 33 unique phyla present in the collared flycatcher GM of which *Proteobacteria* (35.17 %), *Actinobacteria* (31.29 %) and *Firmicutes* (18.10 %) were the most dominant (Fig. 1, SI 1). Other phyla that have also been detected in previous bird studies were *Verrucomicrobia* (1.32%), *Planctomycetes* (4.29%), *Bacteroidetes* (1.60%), *Chlamydiae* (1.29 %), *Tenericutes* (0.98%), *Dependentiae* (0.35%), and *Acidobacteria* (0.95%).

**Figure 1.**
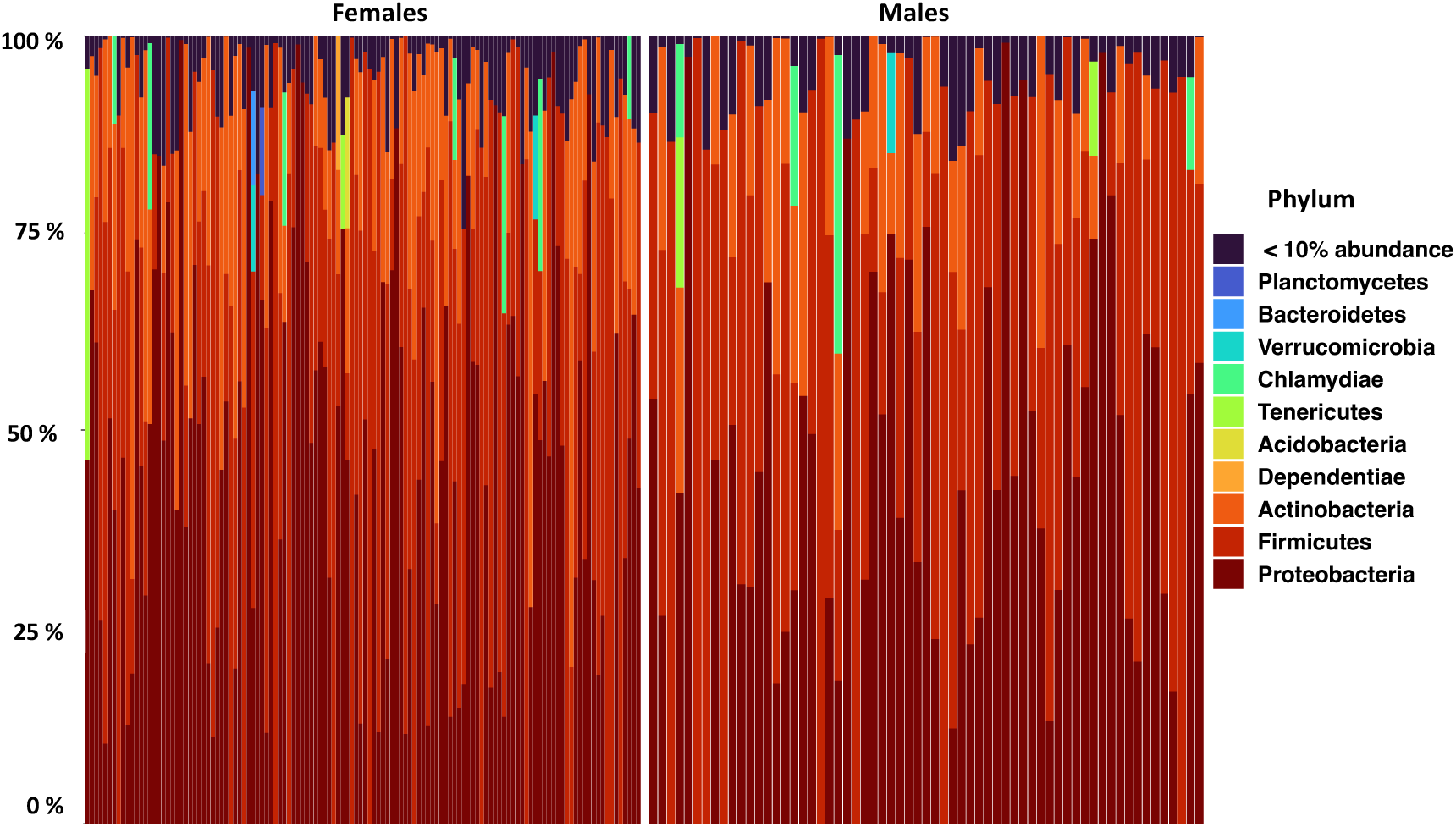
Relative abundance of the most abundant bacterial phyla in the gut microbiome of male and female collared flycatchers. Each bar represents an individual bird. Low abundance phyla are summed up in the “<10% abundance” category for clarity.

### 3.2 Gut microbiome diversity and lifetime reproductive success

Lifetime reproductive success significantly associated with the GM diversity of male collared flycatchers when measured with Shannon: Higher diversity was linked to higher LRS (ANOVA: F_1,38_=6.781, p=0.013, Fig. 2A). This association was not found when all data was used, and there was no association with female LRS (all p=0.344, female p=0.099, Fig. 2B). Chao1 did not significantly associate with LRS in any of the models (P_all_>0.05). Hatch date significantly associated with LRS in all data and females-only data: Earlier hatch date indicated higher LRS (ANOVA_all_: F_1,118_=22.797, p<0.001; ANOVA_females_: F_1,67.601_=13.106, p<0.001). Hatch date did not associate with LRS in males-only data (P=0.336). Body condition and age group were not significant in any of the models (p<0.05). Variation due to breeding area was estimated to be minimal in full dataset (area<0.001, s.d.<0.001, residual variation=11.3, s.d.=3.361) and in male LRS (area<0.001, s.d.<0.001, residual variation=10.06, s.d.=3.172). Variation due to breeding area was a minor in female LRS (area=0.048, s.d.=0.220, residual variation=11.400, s.d.=3.376). Linear mixed effects model estimates are reported in Table 1 and ANOVA tables in Supplementary information 3 (SI 3).

**Figure 2.**
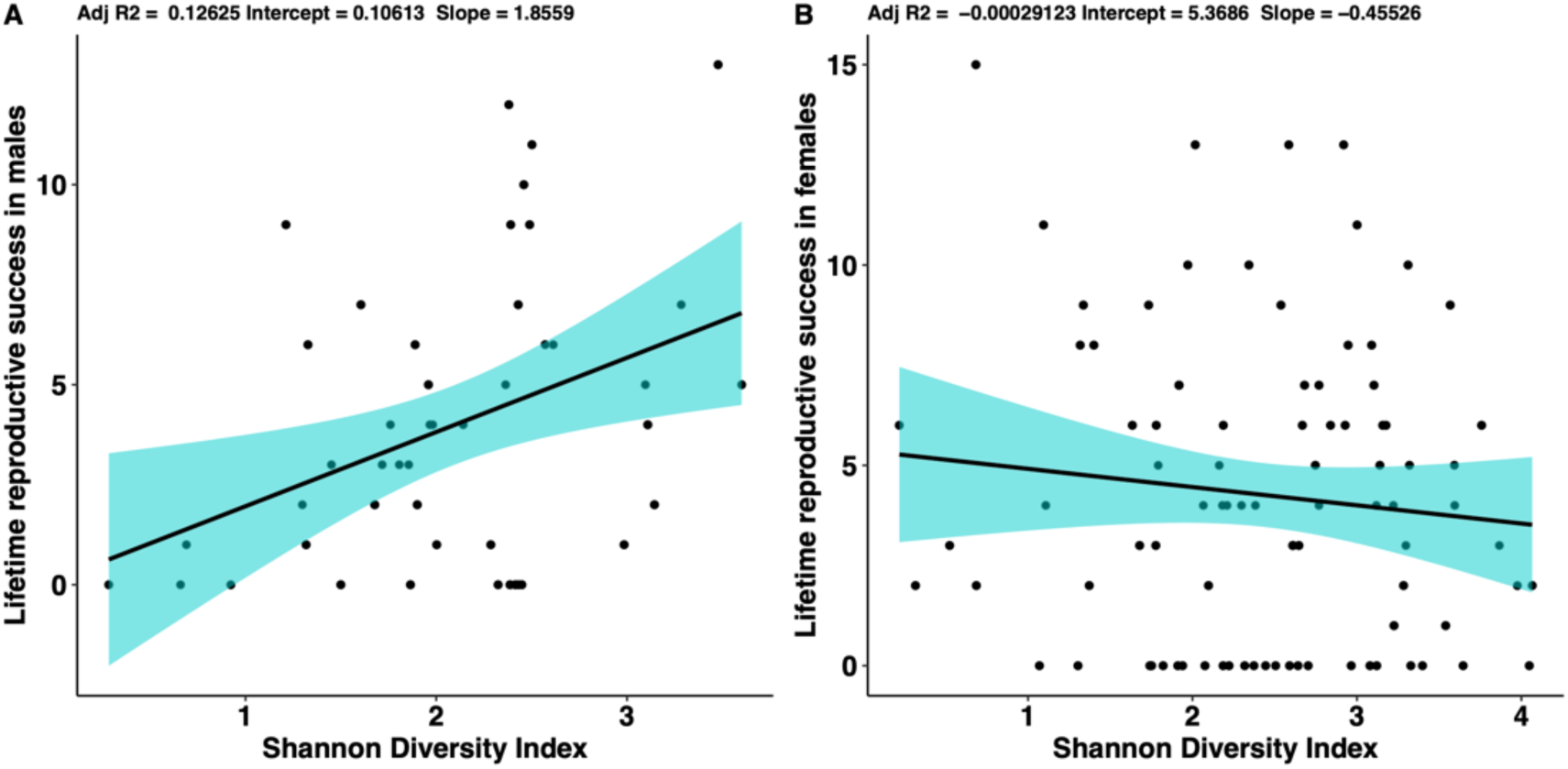
The association between lifetime reproductive success and gut microbiome diversity in A) male and B) female collared flycatchers. The shaded area indicates the s.e.

**Table 1.** Summaries of the linear mixed effects model for A) all collared flycatchers, B) male collared flycatchers, and C) female collared flycatchers when LRS is used as the response variable.

| Predictor | estimate | s.e. | t | P |  | estimate | s.e. | t | P |
| --- | --- | --- | --- | --- | --- | --- | --- | --- | --- |
| **A** |  |  |  |  |  |  |  |  |  |
| Shannon | 0.352 | 0.370 | 0.951 | 0.344 | Chao1 | 0.001 | 0.002 | 0.604 | 0.547 |
| Sex (male) | -0.225 | 0.653 | -0.344 | 0.731 |  | -0.318 | 0.644 | -0.494 | 0.622 |
| Body condition | -0.708 | 0.593 | -1.193 | 0.235 |  | -0.697 | 0.597 | -1.167 | 0.245 |
| Age group 1.5-4 yrs | -0.375 | 0.876 | -0.428 | 0.670 |  | -0.254 | 0.863 | -0.294 | 0.769 |
| Age group 4+ yrs | -1.274 | 1.398 | -0.911 | 0.364 |  | -1.124 | 1.387 | -0.811 | 0.419 |
| Hatch date | -0.346 | 0.072 | -4.775 | **<0.001\*** |  | -0.344 | 0.073 | -4.738 | **<0.001\*** |
| **B** |  |  |  |  |  |  |  |  |  |
| Shannon | 1.982 | 0.761 | 2.604 | **0.013\*** | Chao1 | 0.001 | 0.005 | 0.164 | 0.870 |
| Body condition | 1.076 | 1.081 | 0.996 | 0.326 |  | 0.099 | 1.100 | 0.090 | 0.929 |
| Age group 1.5-4 yrs | 0.523 | 1.594 | 0.328 | 0.745 |  | 1.706 | 1.660 | 1.028 | 0.311 |
| Age group 4+ yrs | -0.142 | 1.899 | -0.075 | 0.941 |  | 0.978 | 2.033 | 0.481 | 0.633 |
| Hatch date | -0.132 | 0.135 | -0.975 | 0.336 |  | -0.245 | 0.140 | -1.743 | 0.089 |
| **C** |  |  |  |  |  |  |  |  |  |
| Shannon | -0.006 | 0.444 | -0.014 | 0.988 | Chao1 | 0.002 | 0.002 | 0.759 | 0.451 |
| Body condition | -1.073 | 0.752 | -1.427 | 0.158 |  | -1.000 | 0.755 | -1.325 | 0.189 |
| Age group 1.5-4 yrs | -0.901 | 1.041 | -0.866 | 0.389 |  | -1.023 | 1.039 | -0.985 | 0.328 |
| Age group 4+ yrs | -2.383 | 2.206 | -1.080 | 0.283 |  | -2.763 | 2.215 | -1.247 | 0.216 |
| Hatch date | -0.342 | 0.094 | -3.620 | **<0.001\*** |  | -0.346 | 0.092 | -3.761 | **<0.001\*** |

\*Sex (female) is used as a reference for sex and age group 1 as a reference for age group.

### 3.3 Gut microbiome diversity and annual reproductive success

GM (Shannon diversity) was significantly positively correlated with ARS when using the full dataset (ANOVA: F_1,115.217_=4.707, p=0.032, Fig. 3C) and when including only males (ANOVA: F_1,35.647_=4.391, p=0.043, Fig. 3A). GM diversity did not significantly associate with the ARS in females (Shannon P=0.099; Chao1 P=0.283, Fig. 3B). Body condition was negatively correlated with ARS when including all data (ANOVA: F_1,117.369_=4.707, p=0.032). Body condition was not significant when sexes were analyzed separately (males p=0.891; females p=0.057). Earlier hatch date correlated with higher ARS in the full and females-only datasets (ANOVA_all_: F_1,81.541_=24.406, p<0.001; ANOVA_females_: F_1,75_=24.128, p<0.001). No association was not found in males-only data (p=0.689). No other explanatory factors correlated with ARS in any of the models (P>0.05). Variation explained by breeding area in males was estimated to be 1.839 (s.d.=1.356, residual variation=3.972, s.d.=1.993), and non-existent in females (area<0.001, s.d.<0.001, residual variation=3.568, s.d.=1.889). In the full dataset this variation was 0.054 (s.d.=0.232, residual variation=3.884, s.d.=1.971). Chao1 did not associate with ARS in any of the models (P_all_>0.05). Linear mixed effects model estimates are reported in Table 2 and ANOVA tables in Supplementary information 3 (SI 3).

**Figure 3.**
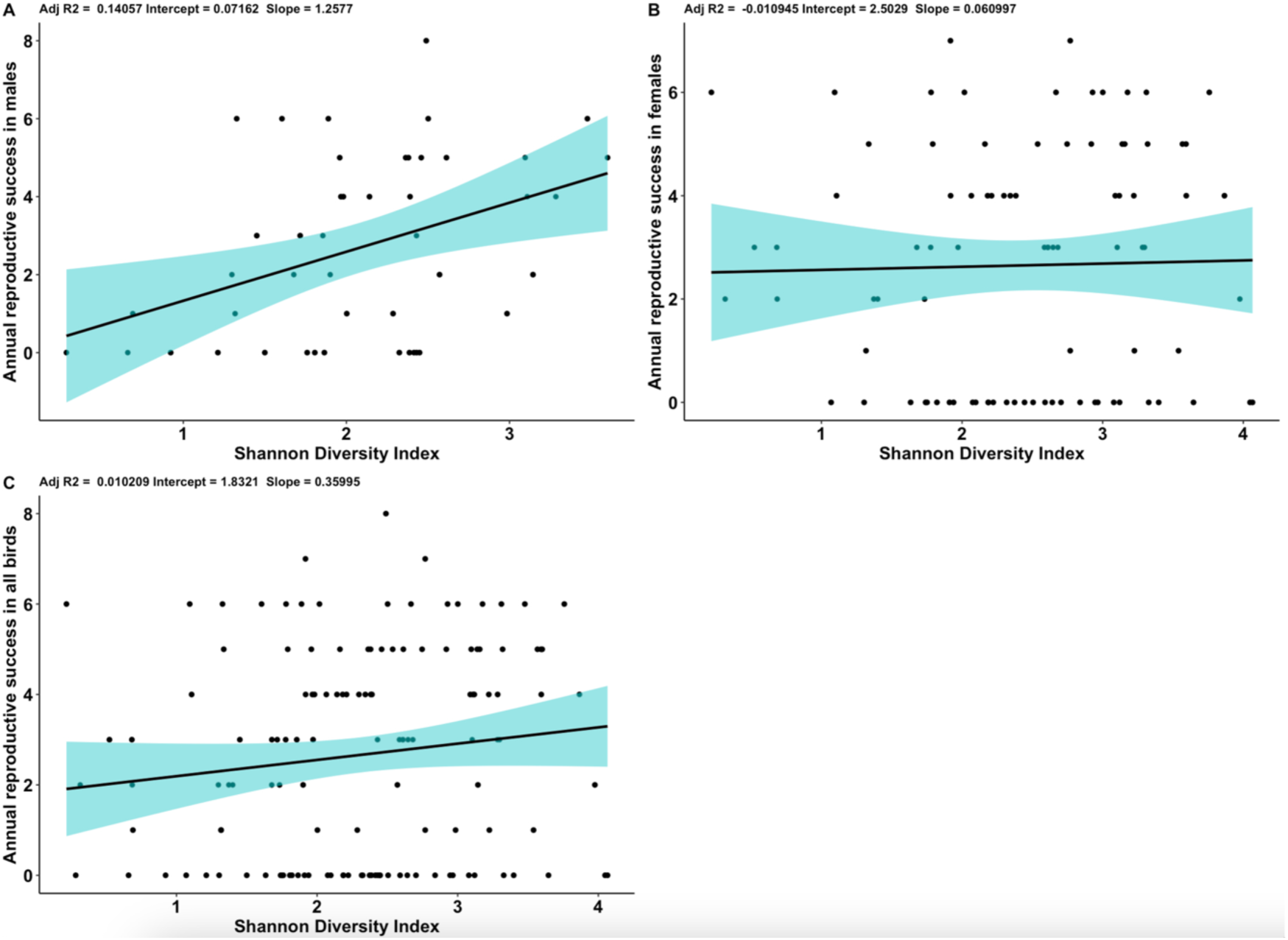
The association of gut microbiome diversity and annual reproductive success in A) male, B) female, and C) all collared flycatchers in 2015. The shaded area indicates the s.e.

**Table 2.**
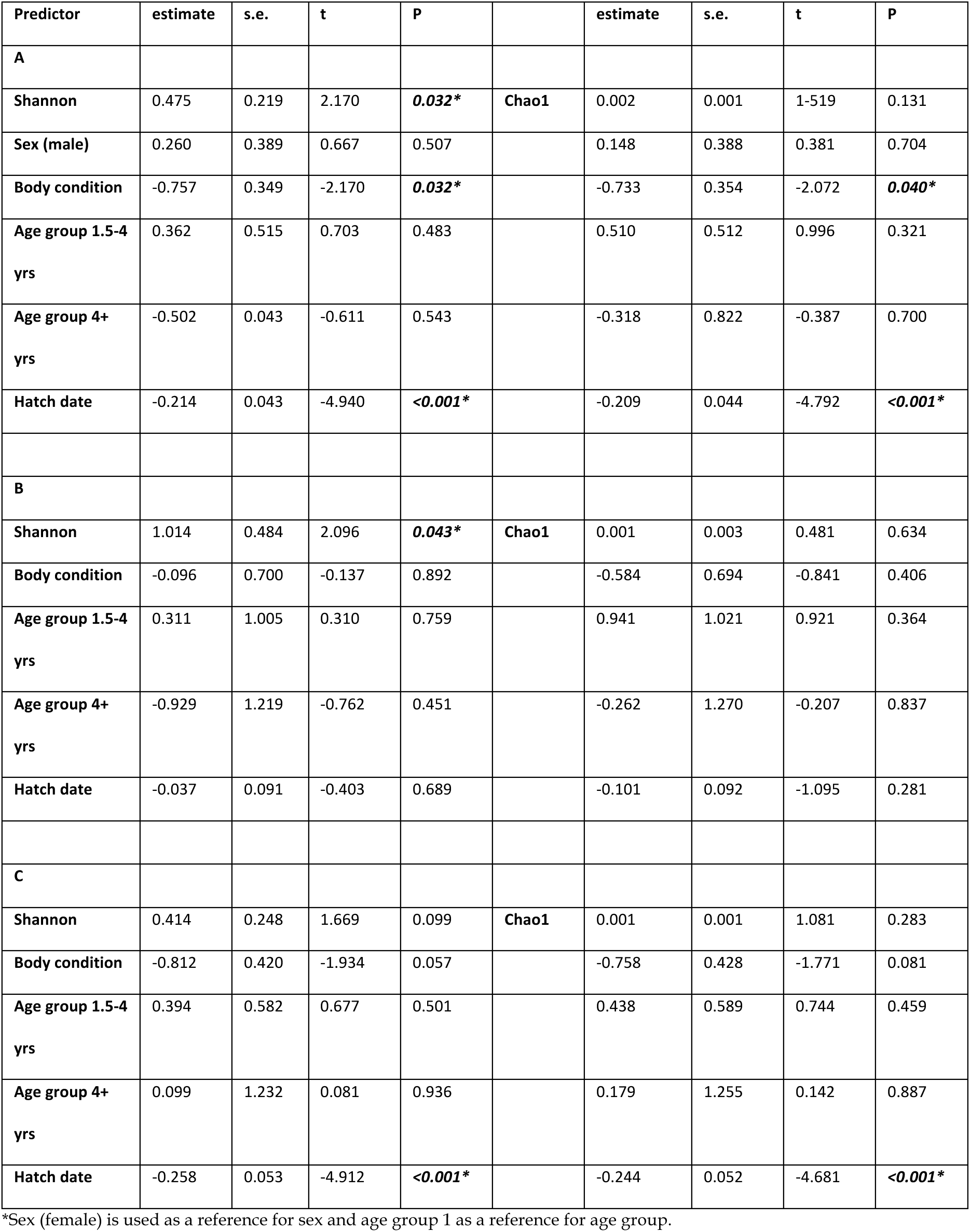
Summaries of the linear mixed effects model for A) all collared flycatchers, B) male collared flycatchers, and C) female collared flycatchers when ARS is used as the response variable.

### 3.4 Gut microbiome diversity and survival to the following breeding season

GM diversity (Shannon and Chao1) did not significantly associate with survival to the following breeding season in any of the models and neither did body condition or age group (P_all,_ P_females_ and P_males_>0.05, SI 4). Sex did not significantly contribute to survival in the model in which both sexes were included (P>0.05, SI 4). Variation explained by breeding area as a random effect was estimated to be non-existent in the full dataset and in males and females (<0.001, s.d.<0.001).

### 3.5 Gut microbiome diversity and environmental and individual characteristics

GM diversity did not associate with body condition, sex, or age group when both sexes were included in the analysis (P_all_>0.05). Variation explained by breeding area was 0.032 (s.d.=0.178, residual variation=0.769, s.d.=0.877). In males-only data, age group was associated with GM diversity (P=0.036): The 1-year-old seemed to have lower diversity than the two older age groups, yet Tukey’s post-hoc test showed that there were no significant differences among groups (P_adj_>0.05). None of the other predicting variables were significant in the males-only model (P_all_>0.05). In females-only data, none of the predicting variables associated with GM diversity (P_all_>0.05). Variation explained by breeding was 0.058 (s.d.=0.241, residual variation=0.785, s.d.=0.886). Model summaries and Tukey’s post-hoc comparisons are reported in Table 3.

**Table 3.**
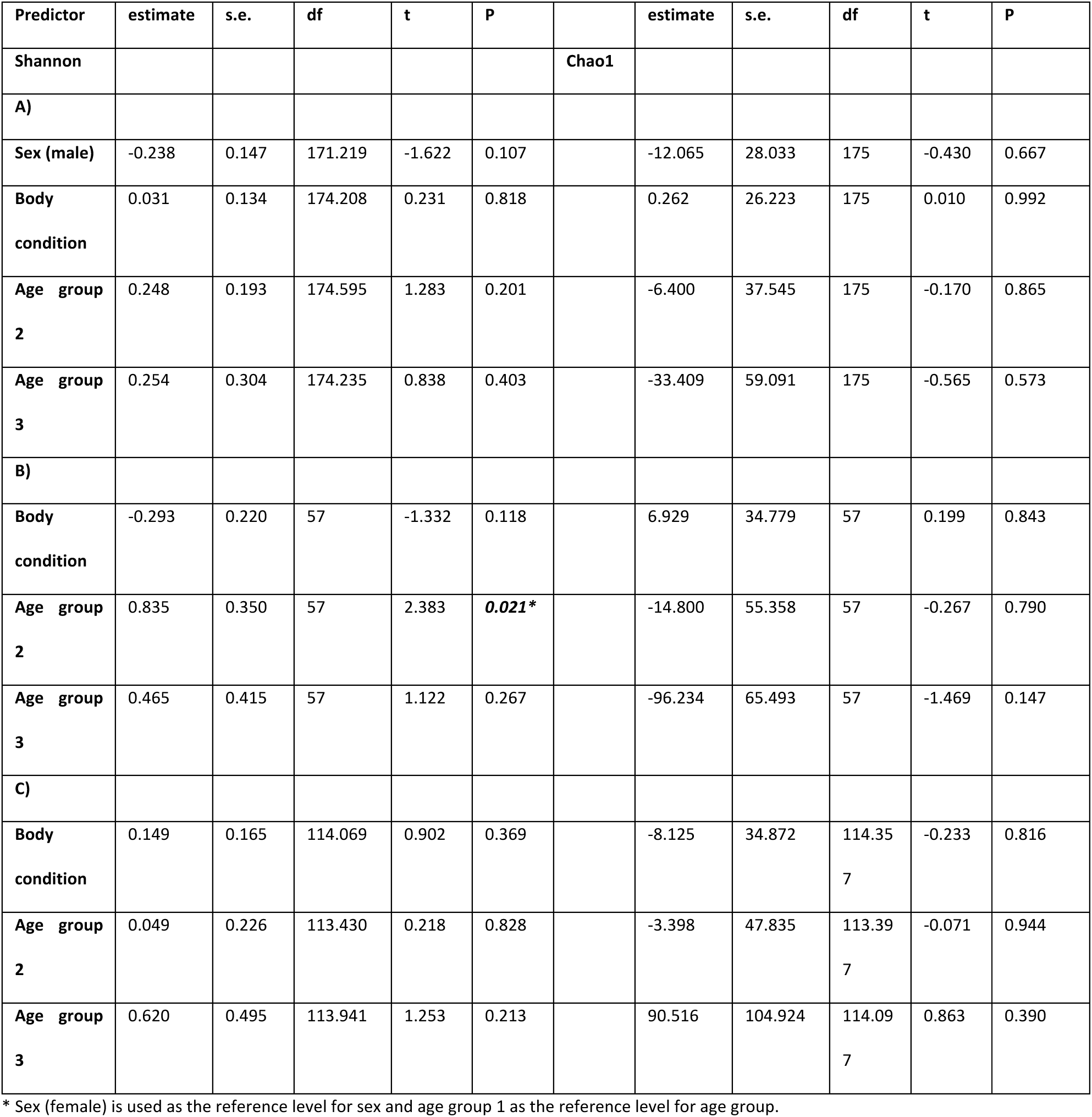
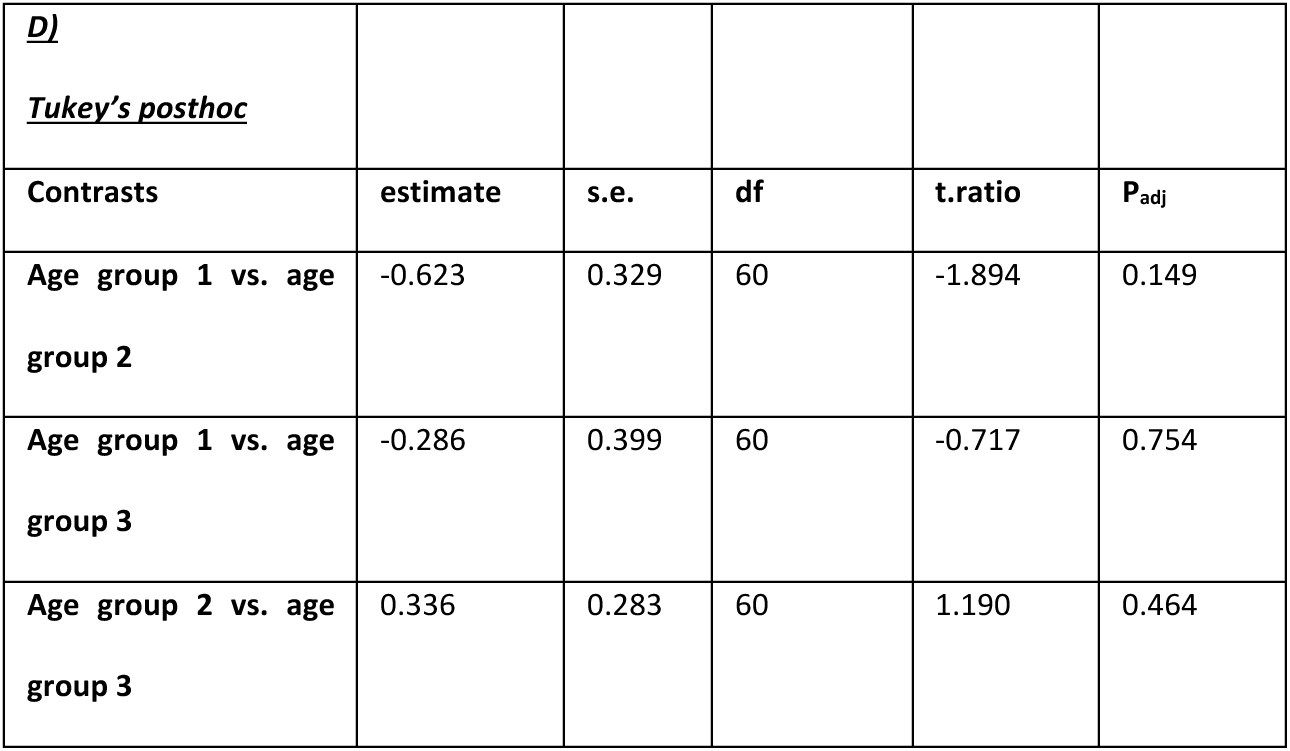
Model summaries of the linear mixed effects model measuring how environmental variables contribute to GM diversity in A) all data, B) males, and C) females. Part D) reports results of Tukey’s post-hoc test for comparisons between male age groups.

### 3.6 Gut microbiome composition

#### 3.6.1 Lifetime reproductive success

LRS did not associate with differences in GM composition in any of the models (PERMANOVA: P>0.05, Fig. 4). Area significantly explained differences in GM composition when both sexes were included in the analysis (PERMANOVA: R^2^=0.083, P=0.025). and in females-only data (PERMANOVA: R^2^=0.132, P=0.012), but not in males-only data (PERMANOVA: P>0.05, SI 6). Age group significantly explained differences in GM composition in males-only data (PERMANOVA: R^2^=0.063, P=0.028), but not in females-only data (PERMANOVA: P>0.05). Area significantly explained differences in GM composition. Body condition and hatch date were not associated with GM composition in any of the models (PERMANOVA: P>0.05). Full model results are reported in Supplementary information 5 (SI 5).

**Figure 4.**
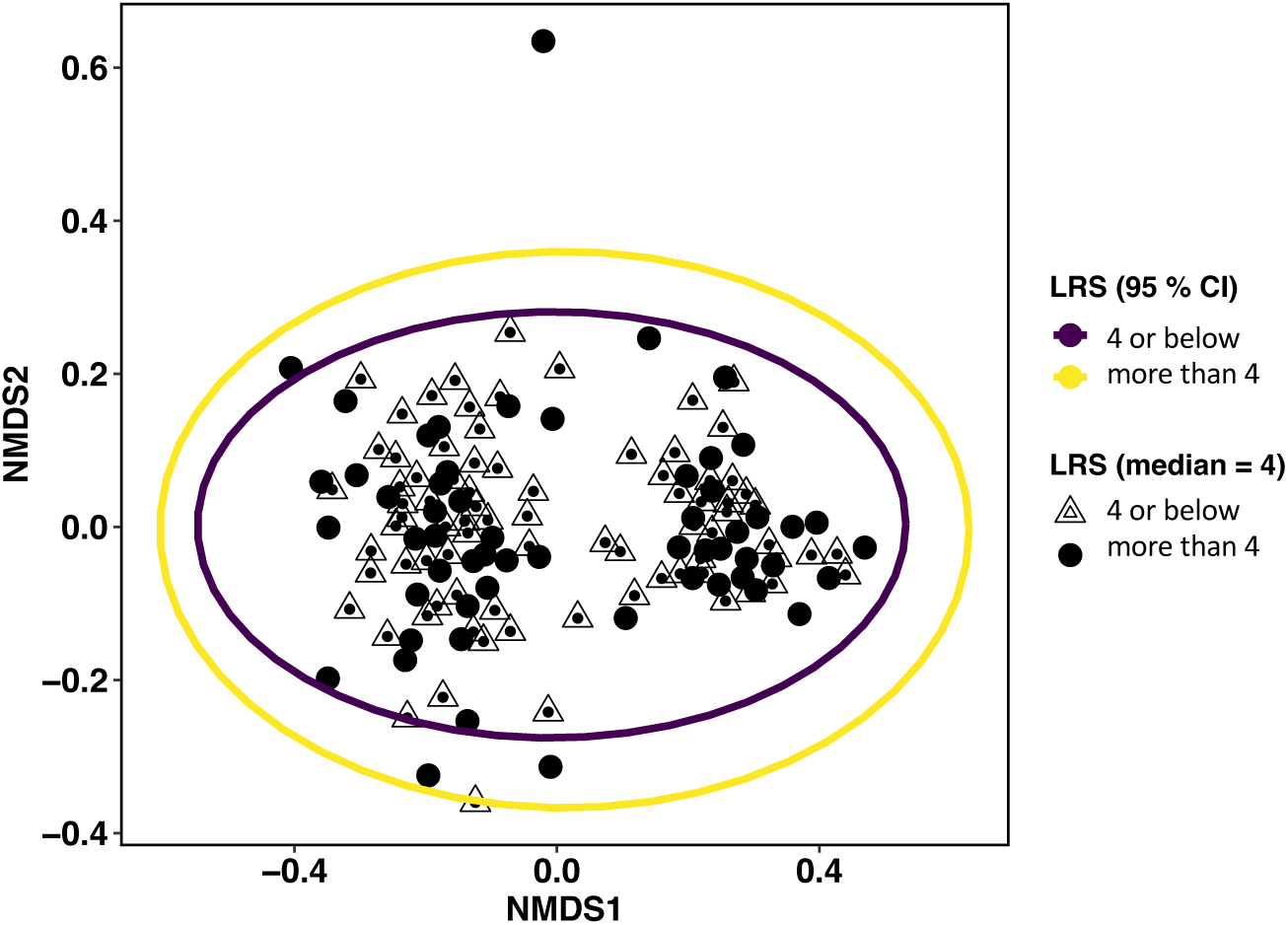
NMDS Bray-Curtis plot visualizing compositional differences in lifetime reproductive success in all data. For visualization, the data was split to two groups: below and above median (median LRS is 4 and mean 4.14) Ellipses represent 95 % confidence intervals.

#### 3.6.2 Annual reproductive success

ARS was not associated with GM composition in any of the models (PERMANOVA: P>0.05). In this model as well, area significantly associated with differences in GM composition when both sexes were included in the model and in females-only, but not males-only, data. Age group significantly associated with differences in GM composition in males-only data but not in the other models. Body condition, hatch date and sex were not significant in any of the models. Full model results are reported in Supplementary information 5 (SI 5).

#### 3.6.3 Survival to the following breeding season

Survival to the following breeding season was not associated with GM composition in any of the models (PERMANOVA: P>0.05, Fig. 5). As in the model above, sex (PERMANOVA: R^2^=0.011, P=0.032) and area (PERMANOVA: R^2^=0.081, P=0.042) were significant contributors to differences in GM composition in the model with both sexes. Area significantly contributed to differences in GM composition in females-only data (PERMANOVA: R^2^=0.125, P=0.016). Body condition and age group were not significant contributors to GM composition in any of the models. Full model results are reported in Supplementary information 5 (SI 5).

**Figure 5.**
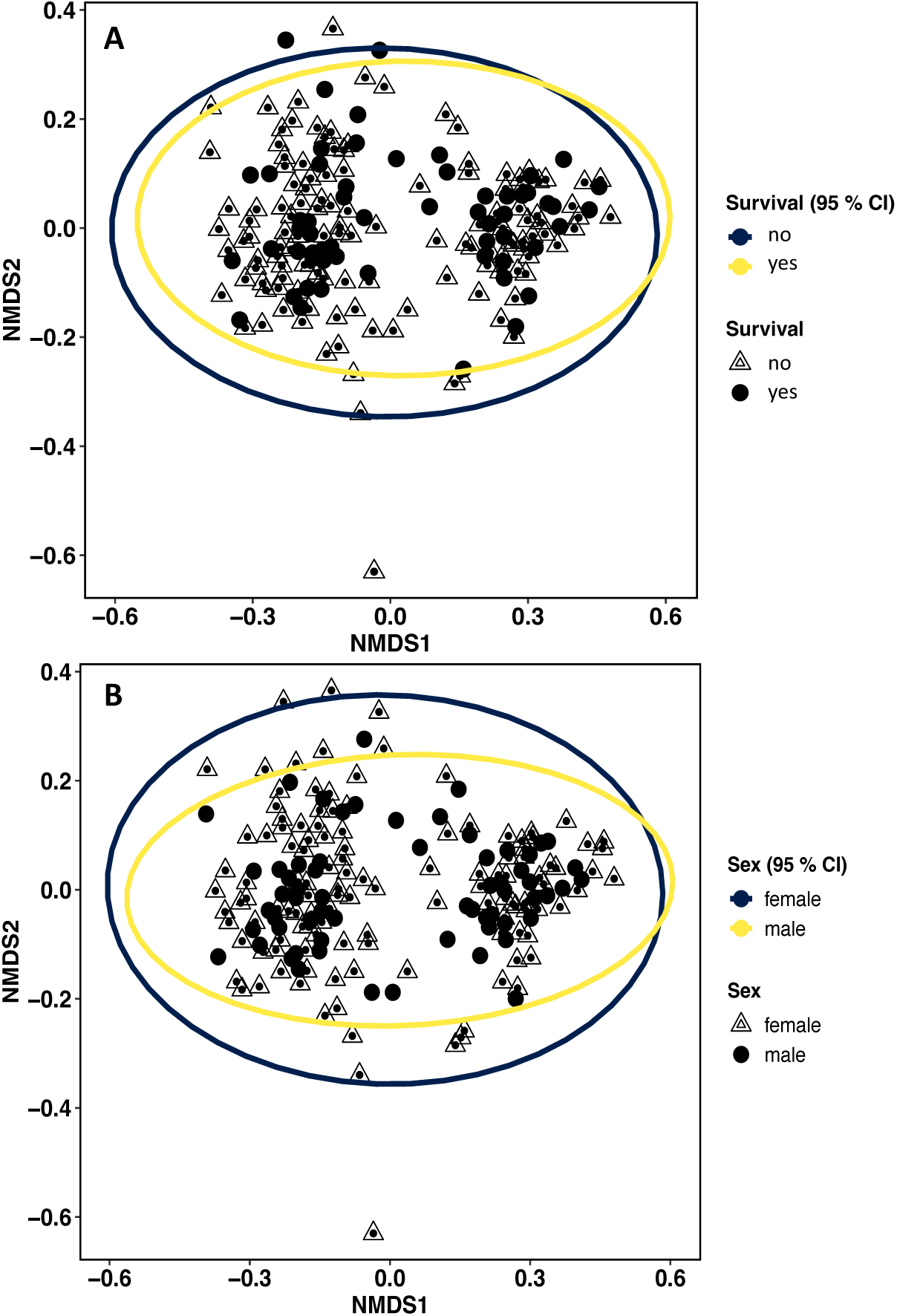
NMDS Bray-Curtis plot visualizing compositional differences in A) survival to the following breeding season and B) sex in individual collared flycatchers. Ellipses represent 95 % confidence intervals.

#### 3.6.4 Differential abundance analysis

There were differentially abundant bacterial taxa in birds that survived to the following breeding season versus those who did not survive (Figure 6, SI 7). When both females and males were included in the analysis, birds that did not survive to the following breeding season had a higher abundance of taxa belonging to the Family *Enterobacteriaceae*. The birds that did not survive also had a higher abundance of the species *Lactobacillus vespulae* and the genera *Corynebacterium* and *Sphingopyxis*. The birds that survived to the following breeding season had a higher abundance of the genera *Dermacoccus*.

**Figure 6.**
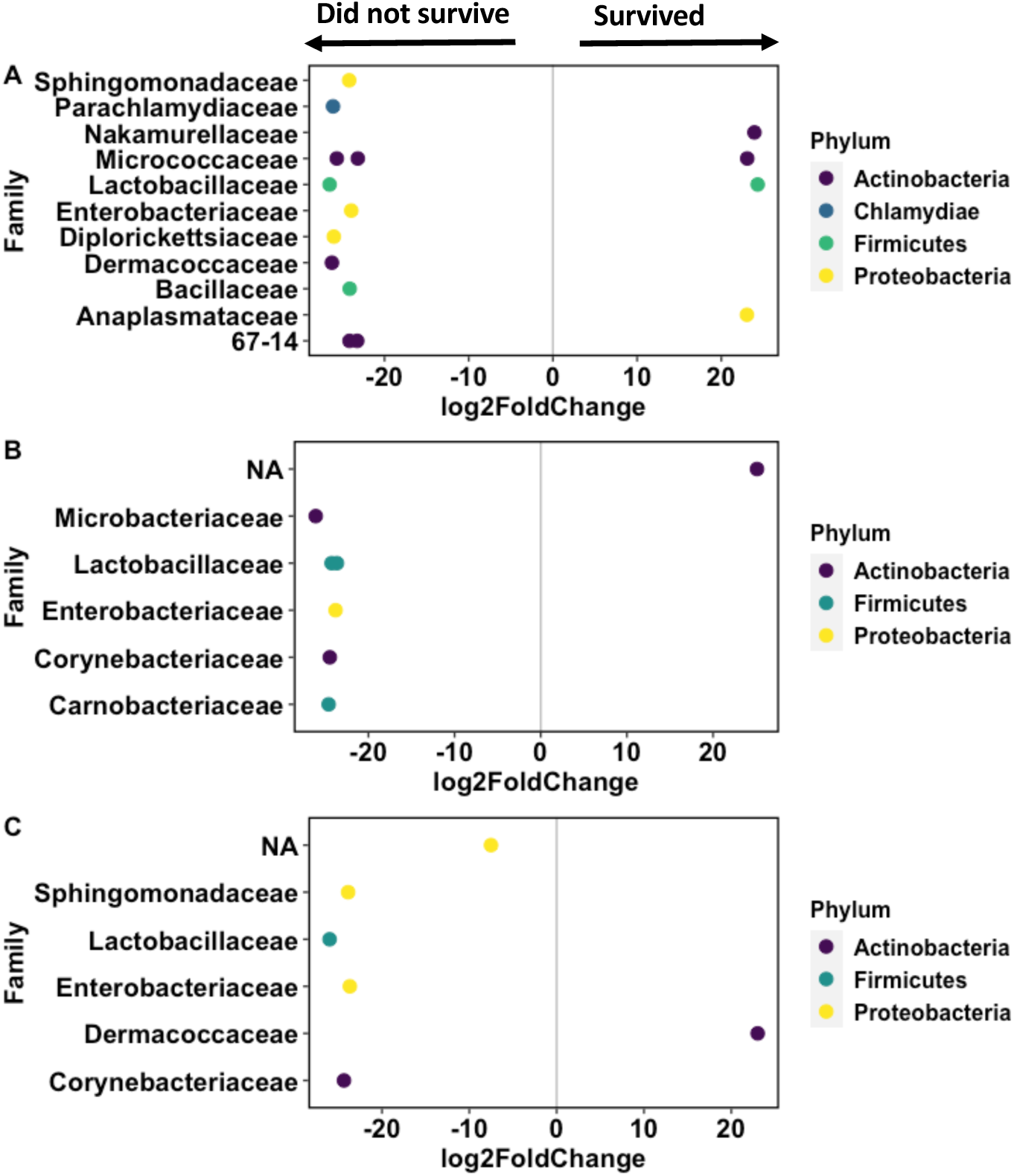
Visualization of the differential abundance analysis of survival to the following breeding season in A) males, B) females, and C) all data. Negative lod2FoldChange represents taxa that are more abundant in birds that did not survive to the following breeding season and positive log2FoldChange represents taxa that are more abundant in birds that survived to the following breeding season.

In males-only data the families *Diplorickettsiaceae*, *Parachlamydiaceae*, *Dermacoccaceae*, *Micrococcaceae* and *Enterobacteriaceae* were more abundant in birds that did not survive to the following breeding season compared to the birds that survived. Also, the genus *Sphingomonas* and the species *Bacillus simplex* were more abundant in birds that did not survive. In birds that survived to the following breeding season the genera *Nakamurella*, *Glutamicibacter* and *Wolbachia* were more abundant compared to the birds that did not survive. The species *L. vespulae* was found in both birds that survived and birds that did not survive to the following breeding season, but its abundance varied little between the groups.

In females-only data the families *Microbacteriaceae* and *Enterobacteriaceae* were more abundant in the birds that did not survive to the following breeding season. Also, the species *L. vespulae* and the genera *Carnobacterium* and *Corynebacterium* were more abundant in the birds that did not survive. In birds that survived to the following breeding season, an unidentified bacterial order belonging to the class *Acidimicrobiia* was more abundant than in birds that did not survive.

## 4 DISCUSSION

We investigated whether the collared flycatcher GM diversity and composition associate with individual LRS, ARS and survival to the breeding season following the sampling year. Our results showed that GM diversity (Shannon, but not Chao1) was positively associated with ARS, and sex-specific investigation revealed that this positive association was significant in males. Furthermore, GM diversity was positively associated with LRS in males. We also found that hatch date was associated with both ARS and LRS and that this association was specific to female collared flycatchers. In our study, we found no significant association between ARS, survival to the following breeding season or LRS when Chao1 was used as diversity metric. Concerning GM composition, LRS, ARS or survival were not associated with GM composition. Yet there were differentially abundant taxa between the birds that survived to the following breeding season and the birds that did not survive. Finally, sex significantly associated with differences in GM composition but not diversity. Additionally, in males GM composition and diversity differed across age groups, while in females, birds originating from different areas showed differences in GM composition and diversity. This is the first study to characterize the collared flycatcher GM and investigate the association between the GM and reproductive success in wild birds. It emphasizes that future research should study whether there is a causal relationship between the gut microbiome and reproductive success or if the correlation we found in this study is a result of some unknown third factor. Further studies could utilize disruptive methods such as exposure to antibiotics to investigate whether variation in the gut microbiome affects reproductive success (Rosengaus et al., 2011).

The collared flycatcher GM is dominated by the phyla *Proteobacteria*, *Actinobacteria*, and *Firmicutes*, which is comparable with previous studies done with small passerine birds (Bodawatta et al., 2021; Drobniak et al., 2022; Hird et al., 2015; Somers et al., 2023; Teyssier et al., 2018). Furthermore, we observed high individual variation among the fecal samples, which was expected (Kropáčková et al., 2017). As migratory birds, collared flycatcher adults are exposed to varying environments, which may explain variation in their GM (Grond et al., 2018). Because collared flycatcher GM has not been sequenced before, we cannot compare our results to previous studies done with same species. However, there is GM data from the same research area in which blue tit GM has been characterized: The blue tits harbored the same core phyla *Proteobacteria*, *Firmicutes* and *Actinobacteria* as did the collared flycatchers in our study (Drobniak et al., 2022). This can be a result of both environmental factors and life-history.

### 4.1 Gut microbiome and reproductive success

GM diversity (as measured with Shannon), but not composition, associated with both LRS and ARS, and this association was most likely explained by the male birds in our study: There was a strong positive correlation between high GM diversity and LRS / ARS in male but not female collared flycatchers. Generally, individuals that are in better condition are more likely to survive and produce high quality offspring (Blomqvist et al., 1997; Jensen et al., 2004; Pigeault et al., 2020; Wendeln and Becker, 1999) and similar pattern has been observed in collared flycatchers as well (Szöllősi et al., 2009). However, in this study we found that there is a slight negative correlation between body condition and ARS. High GM diversity is often linked to better individual quality because a more diverse GM may often have functionally similar taxa, which makes the GM more stable (as reviewed in Lozupone et al., 2012). Therefore, it is possible that the high GM diversity in especially male collared flycatchers was a factor underlying individual quality and reproductive success in our study as well. However, we did not find a correlation between body condition index and GM diversity, yet other predictors of individual quality may explain the association. Also, previous studies are inconclusive on whether the GM directly contributes to body condition (e.g., Phillips et al., 2018; Potti et al., 2002; Teyssier et al., 2018; Videvall et al., 2019; Worsley et al., 2021).

Given that the positive correlation between GM diversity and LRS / ARS was found in males, but not in females, this indicates that there may be sex-specific associations between GM diversity and reproductive success. We may speculate that a potential link, specific to males, may be via the association between GM, testosterone and breeding success: increased testosterone levels have been linked to high GM diversity in mammals (Markle et al., 2013; Shin et al., 2019) and there is some evidence that the GM can regulate testosterone production even though the mechanisms underlying this regulation are still unclear (Colldén et al., 2019). Interestingly, higher testosterone levels were associated with increased cloacal bacterial diversity (which partially resembles GM) in male rufous-collared sparrows *Zonotrichia capensis* (Escallón et al., 2017). High testosterone level affects behavior and can increase aggressiveness, which can improve both parental quality and securing a higher quality territory and therefore, improve breeding success (Szász et al., 2019). This may suggest that the association between high reproductive success and high GM diversity may be linked to testosterone. Additionally, higher testosterone levels can also increase EPC and EPP behaviour in males. Males that copulate with “extra” females have a chance of producing more offspring than the males with lower testosterone levels (Wingfield 1984, Raouf et al., 1997, Grear et al., 2009). EPC also subjects the male to more social encounters and cloacal inoculation of bacteria during copulation (Escallón et al., 2019). Few studies have found inconclusive results regarding EPC and EPP’s role in bird gut microbiomes (Kreisinger et al., 2015, Escallón et al., 2019, Prüter et al., 2023). EPC and EPP were not investigated in this study but should be taken into account in future studies.

Interestingly, in a study with blue tits, parental quality was especially important in male birds in terms of reproductive success but not female birds (Przybylo et al., 2001) and this may explain the difference between sexes in our study as well. Furthermore, collared flycatchers are migratory birds and migrate annually between their breeding and wintering grounds. Male collared flycatchers migrate to their breeding grounds before the females and females choose the males based on the territory quality (Gustafsson, 1985). Therefore, it could be that the male GM is more strongly associated with reproductive success via territory quality and breeding behavior. Finally, we also found a significant association between hatch date and ARS and LRS of females, which was expected as it has been well reported in multiple previous studies. Females that breed earlier and have their young hatch earlier are more successful in rearing their young and thus, have higher reproductive success (e.g., Halupka et al., 2021; Siikamäki, 1998; Verboven and Visser, 1998; Verhulst et al., 1995).

In contrast to GM diversity, we did not find association between GM composition and ARS or LRS. These findings contrast with the evidence so far: There is increasing evidence of a link between hormone production, reproduction and GM composition even though most studies are still purely correlative (Neuman et al., 2015; Williams et al., 2020) and data from birds is not available. In female eastern black rhinos GM composition and higher abundance of specific bacterial genera associated reproductive functioning (Antwis et al., 2019) and similar association was found in the southern white rhinoceros *Ceratotherium simum simum* (Burnham et al., 2023) and Phayre’s leaf monkeys *Trachypithecus phayrei crepusculus* (Mallott et al., 2020). In a study with captive Asian *Elephas maximus* or African *Loxodonta africana* elephants the GM correlated with stress hormones but not reproductive hormones (Keady et al., 2021). Furthermore, significant changes in crested ibis GM composition and increased *Proteobacteria* abundance correlated with reproductive dysfunction (Ran et al., 2021). However, the causal link between GM composition and reproductive success has not been found in vertebrates so far.

### 4.2 Gut microbiome and survival to the following breeding season

We found no association between the GM diversity or composition and survival to the following breeding season. This contrasted with a previous study in which GM composition associated with survival to the following breeding season in wild Seychelles’ warblers (Worsley et al., 2021). In our study, we use adult birds that have survived one or more migrations and have successfully returned to breed. Quite often the first migration can be the deadliest for juvenile birds as they are inexperienced, start migration later than their adult counterparts, fail to secure a breeding territory or die during migration (McKim-Louder et al., 2013; Oppel et al., 2015). The lack of association between GM diversity and survival could result from selective disappearance, i.e. the sampled birds being essentially more fit to begin with than their counterparts that did not survive. We also must bear in mind that there is a small possibility that the birds that we categorized as “not survived” could also have relocated to some other breeding area that we did not monitor (unlikely as collared flycatchers have a high return rate: Gustafsson, 1986; Part and Gustafsson, 1989).

However, there were differentially abundant taxa between the birds that survived to the following breeding season and the birds that did not survive. Unfortunately, many bacterial taxa are still unknown and their role in the GM and host health need to be studied (Dias et al., 2020). Here, we focus on discussing the bacterial taxa that are known of their influence on host health. In birds that did not survive, we observed a higher abundance of the species *Lactobacillus vespulae*, which belongs to the genus *Lactobacillus* and is known for its benefits for host health. The result is curious as usually *Lactobacillus* are connected to better health and improve survival (Davidson et al., 2021; Reid and Burton, 2002; Worsley et al., 2021). *L. vespulae* was first isolated from the gut of the queen wasp *Vespula vulgaris* (Hoang et al., 2015). Possible explanations for this are that 1) the birds with a higher abundance of this *Lactobacillus* sp. have either nested in a nest box in which wasps have previously nested (Broughton et al., 2015) or fed on wasps (Mäntylä et al., 2015). The birds that did not survive to the following breeding season also had a higher abundance of the genera *Corynebacterium*, *Sphingopyxis* and *Dermacoccus*. *Corynebacterium* have been observed to cause infections especially in immunocompromised human patients (Bernard, 2012). In birds, a study with magellanic penguin *Spheniscus magellanicus* chicks showed that *Corynebacterium* can decrease growth by altering gut metabolism and thus, influence survival (Potti et al., 2002).

When we ran the models separately for each sex, we found that the male collared flycatchers that did not survive had a higher abundance of the Families *Enterobacteriaceae*, *Diplorickettsiaceae*, *Parachlamydiaceae*, the genus *Sphingomonas,* and the species *Bacillus simplex*, which are all potential pathogens. For example, *Enterobacteriaceae* has been suspected to contribute to the bloom of harmful taxa in the gut and enhance inflammatory responses that can reflect to individual health (Baldelli et al., 2021), and *Parachlamydiaceae* is known to cause infections in captive birds (Frutos et al., 2015; You et al., 2019). The genus *Sphingomonas* can cause severe infections in individuals that are already immunocompromised (Ryan and Adley, 2010; White et al., 1996), and pathogenic *Bacillus* strains have been found on birds previously and they can be transmitted via e.g., feather contact (Miskiewicz et al., 2018). In males, testosterone levels are increased during the breeding season and the higher concentration could decrease immune system functioning (Immunocompetence Handicap Hypothesis) (Rantala et al., 2012; Roberts et al., 2004) (but see: Nowak et al., 2018) and thus, may explain the high number of potentially pathogenic taxa. Conversely, in the birds that survived, we found a higher abundance of the genus *Wolbachia* that can potentially improve host health (Serbus et al., 2008). *Wolbachia* usually infects insect and nematodes and can live as a symbiont in the host gut and improve host fitness by limiting other potential pathogens and increase antiviral protection (Hedges et al., 2008). In birds *Wolbachia* have been detected in quill mites (Glowska et al., 2015) but the possible effects on health have not been discussed. In female collared flycatchers, the family *Enterobacteriaceae*, the genus *Corynebacterium* and the species *L. vespulae* were also more abundant in birds that did not survive. However, specific to female collared flycatchers only, we found that the family *Microbacteriaceae* and the genus *Carnobacterium* were more abundant in birds that did not survive than in the birds that survived, but the function of these taxa in the bird gut is not known.

### 4.3 Environmental and intrinsic factors contributing to GM variation in collared flycatchers

As data on the environmental and intrinsic factors explaining GM variation in collared flycatchers is not available, we also report and briefly discuss key factors that contributed to GM variation. Sex explained 1.1 % and breeding area (plot) 8.3 % of compositional differences in the GM. Furthermore, age group explained 6.3 % of compositional differences in male collared flycatcher GM and was associated with GM diversity, but this was not found in females. Breeding area explained 13.2 % of compositional differences and explained part of the variation in GM diversity in female collared flycatchers, and but not in males. The between-sex variation (although very minor) in GM composition could be a result of sex-specific differences in physiology and behavior as they have been observed in many species including mammals and birds (Liu et al., 2020; Markle et al., 2013; Ren et al., 2017; Worsley et al., 2021; Yan et al., 2022) (but see Drobniak et al., 2022a; Kohl et al., 2017). For example, in thick-billed murres *Uria lomvia* GM composition varied between sexes because of sex-specific behavioral feeding differences: (Góngora et al., 2021). Similar sex-specific differences in GM composition that could result from dietary preferences have also been identified in the great bustard *Otis tarda* (Liu et al., 2020). It is possible that the minor sex-specific variation in GM composition that we observed here connects to some physiological differences between sexes such as hormone system functioning (Escallón et al., 2019). Further investigation is needed to understand whether this sex-specific variation contributes to individual physiology and even, reproductive success.

GM composition can be habitat specific (Bletz et al., 2016; Drobniak et al., 2022). Previous studies have found breeding site quality to be one of the drivers of differences in the GM composition of shorebirds (Grond et al., 2019) and greater flamingos *Phoenicopterus roseus* (Gillingham et al., 2019). In our study, breeding area is likely to influence the GM composition indirectly via various environmental factors, especially food abundance and quality, that are known to alter the GM composition and diversity (Bodawatta et al., 2021; Góngora et al., 2021; Loo et al., 2019). Moreover, the association between age group and GM composition can potentially be a result of aging (Bosco and Noti, 2021; Jia et al., 2018) or potentially also reflect territory or even diet quality. For example, in rhesus macaques *Macaca mulatta* different age groups showed differences in GM composition but not in GM diversity (Adriansjach et al., 2020). In birds, age has been found to associate with the GM composition of for example adult Seychelles’ warblers (Worsley et al., 2023) and both chick and adult black-legged kittiwakes *Rissa tridactyla* (Van Dongen et al., 2013).

## 5 CONCLUSIONS

This is the first study to characterize the collared flycatcher GM and to investigate how the GM associates with lifetime reproductive success in wild birds. The results bring much needed knowledge about the GM and Darwinian fitness in a wild population. We show that the GM diversity associates with reproductive success in especially male collared flycatchers. We did not find any association between GM composition and reproductive success or survival, which was curious and needs more investigation. However, we found a higher abundance of pathogenic taxa in birds that did not survive to the following breeding season than in the birds that survived. This suggests that the presence pathogenic bacterial taxa can influence host health and overall fitness. Even though these results are correlational, they provide important knowledge of the collared flycatcher GM and how it associates with reproductive success in wild birds. Future studies should focus on understanding the function of avian GMs and how the GM may influence individual endocrine system functioning, parental quality, and fitness. Furthermore, it should be investigated whether the GM is driving differences in survival to the following breeding season and LRS or vice versa.

## Supporting information

Supplemental files for manuscript

## AUTHOR CONTRIBUTIONS

Martta Liukkonen processed the samples and resulting sequence data, analyzed the data and wrote the manuscript. Lars Gustafsson collected the samples, organized, and performed fieldwork and field data. Kirsten Grond helped to write the bioinformatics script. Suvi Ruuskanen assisted with sample processing, data analysis and manuscript writing. All authors read and commented on the final manuscript.

## ACKNOWLEDGEMENTS

We would like to thank Emil Aaltonen Foundation for funding this research, CSC – IT center for science for computing power, and Dr. Charli Davies for helping with the bioinformatics script.

## ETHICAL STATEMENT

For studying wild birds, we had permits from the Swedish ringing center – license 471. Birds were not harmed during the study.

## DATA AVAILABILITY

Sequences will be archived in University of Jyväskylä’s NextCloud and NCBI Sequence Read Archives (SRA). QIIME2 and R scripts will be available in GitHub (https://github.com/marttal/flycatchers) and on request from the corresponding author.

