## Supplemental files for manuscript for "Gut microbiome diversity associates with estimated lifetime and annual reproductive success in male but not female collared flycatchers"

### SI 1. Sequencing results

| | Number of samples | Number of taxa | Min.reads | Max.reads | Total reads | Mean reads | Median reads |
|---|---|---|---|---|---|---|---|
| **Unrarefied** | 187 | 22 632 | 260 | 853 017 | 33 726 887 | 170 337.813 | 79 947.5 |
| **Rarefied (2000 taxa)** | 185 | 6623 | 2000 | 2000 | 392 000 | 2000 | 2000 |

### SI 2: Taxa prevalence at Phylum level across all samples

The prevalence of taxa across all samples in the observed 22 632 ASVs in the unrarefied dataset. For clarity, the taxa are plotted at Phylum level. Each dot represents an individual ASV within a phylum.

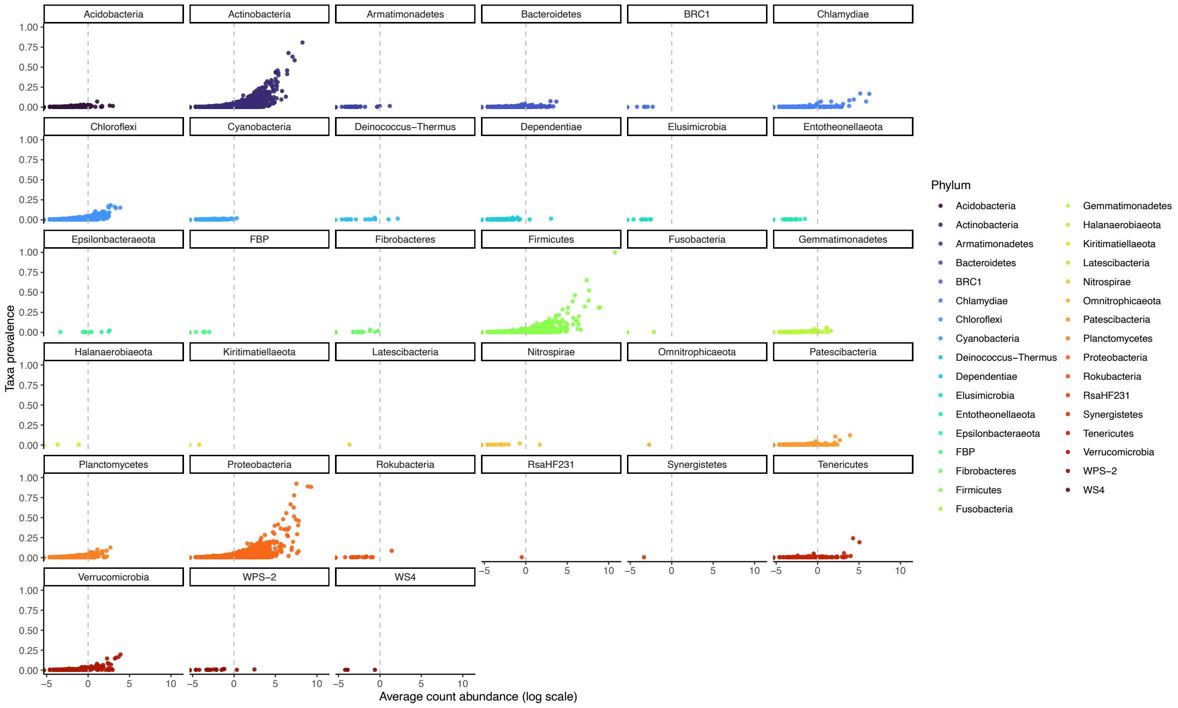

### SI 3: Results of the linear mixed effects model measuring the association between gut microbiome diversity and LRS / ARS

Table 1. The ANOVA type III (Satterthwaite's method) table of the linear mixed effects models on the association between gut microbiome diversity (Shannon and Chao1) and LRS in A) all collared flycatchers, B) in male collared flycatchers, C) in female collared flycatchers, and ARS in D) all collared flycatchers, E) in male collared flycatchers, and in F) female collared flycatchers.

| **Response** | **SumSq** | **df** | **ddf** | **F** | **P** | | **SumSq** | **df** | **ddf** | **F** | **P** |
|---|---|---|---|---|---|---|---|---|---|---|---|
| **A)** | | | | | | | | | | | |
| **Shannon** | 10.209 | 1 | 118 | 0.903 | 0.344 | **Chao1** | 4.135 | 1 | 118 | 0.364 | 0.547 |
| **Sex** | 1.338 | 1 | 118 | 0.119 | 0.731 | **Sex** | 2.772 | 1 | 118 | 0.244 | 0.622 |
| **Body condition** | 16.064 | 1 | 118 | 1.422 | 0.235 | **Body condition** | 15.463 | 1 | 118 | 0.339 | 0.713 |
| **Age group** | 9.420 | 2 | 118 | 0.417 | 0.660 | **Age group** | 7.686 | 2 | 118 | 0.339 | 0.713 |
| **Hatch date** | 257.496 | 1 | 118 | 22.797 | *<0.001** | **Hatch date** | 254.703 | 1 | 118 | 22.448 | *<0.001** |
| **B)** | | | | | | | | | | | |
| **Shannon** | 68.224 | 1 | 38 | 6.781 | *0.013** | **Chao1** | 0.320 | 1 | 38 | 0.027 | 0.870 |
| **Body condition** | 9.977 | 1 | 38 | 0.992 | 0.326 | **Body condition** | 0.096 | 1 | 38 | 0.008 | 0.929 |
| **Age group** | 2.667 | 2 | 38 | 0.133 | 0.876 | **Age group** | 13.208 | 2 | 38 | 0.557 | 0.577 |
| **Hatch date** | 9.570 | 1 | 38 | 0.951 | 0.336 | **Hatch date** | 35.992 | 1 | 38 | 3.038 | 0.089 |
| **C)** | | | | | | | | | | | |
| **Shannon** | 0.002 | 1 | 73.340 | 0.0002 | 0.998 | **Chao1** | 6.510 | 1 | 75.000 | 0.575 | 0.451 |
| **Body condition** | 23.209 | 1 | 74.858 | 2.036 | 0.158 | **Body condition** | 19.856 | 1 | 74.964 | 1.755 | 0.189 |
| **Age group** | 15.722 | 2 | 74.485 | 0.690 | 0.505 | **Age group** | 20.563 | 2 | 74.551 | 0.909 | 0.408 |
| **Hatch date** | 149.406 | 1 | 67.601 | 13.106 | *<0.001** | **Hatch date** | <0.001 | 1 | 70.379 | 14.144 | *<0.001** |
| **D)** | | | | | | | | | | | |
| **Shannon** | 18.281 | 1 | 115.217 | 4.707 | *0.032** | **Chao1** | 9.136 | 1 | 117.184 | 2.309 | 0.131 |
| **Sex** | 1.726 | 1 | 81.895 | 0.444 | 0.507 | **Sex** | 0.574 | 1 | 83.527 | 0.145 | 0.704 |
| **Body condition** | 18.282 | 1 | 117.369 | 4.707 | *0.032** | **Body condition** | 16.990 | 1 | 117.658 | 4.293 | *0.040** |
| **Age group** | 7.108 | 2 | 116.147 | 0.915 | 0.403 | **Age group** | 8.295 | 2 | 116.467 | 1.048 | 0.354 |
| **Hatch date** | 94.791 | 1 | 81.541 | 24.406 | *<0.001** | **Hatch date** | 90.874 | 1 | 89.012 | 22.965 | *<0.001** |
| **E)** | | | | | | | | | | | |
| **Shannon** | 17.442 | 1 | 35.647 | 4.391 | *0.043** | **Chao1** | 1.028 | 1 | 35.489 | 0.231 | 0.634 |
| **Body condition** | 0.075 | 1 | 36.511 | 0.019 | 0.891 | **Body condition** | 3.151 | 1 | 36.718 | 0.708 | 0.406 |
| **Age group** | 7.025 | 2 | 36.270 | 0.884 | 0.422 | **Age group** | 8.593 | 2 | 36.115 | 0.965 | 0.391 |

| | | | | | | | | | | | |
|---|---|---|---|---|---|---|---|---|---|---|---|
| **Hatch date** | 0.647 | 1 | 36.325 | 0.163 | 0.689 | **Hatch date** | 5.337 | 1 | 36.523 | 1.199 | 0.281 |
| **F)** | | | | | | | | | | | |
| **Shannon** | 9.944 | 1 | 75 | 2.787 | 0.099 | **Chao1** | 4.256 | 1 | 75 | 1.168 | 0.283 |
| **Body condition** | 13.343 | 1 | 75 | 3.739 | 0.057 | **Body condition** | 1.430 | 1 | 75 | 3.137 | 0.081 |
| **Age group** | 1.805 | 2 | 75 | 0.253 | 0.777 | **Age group** | 2.140 | 2 | 75 | 0.294 | 0.746 |
| **Hatch date** | 86.099 | 1 | 75 | 24.128 | *<0.001** | **Hatch date** | 79.837 | 1 | 75 | 21.908 | *<0.001** |

* Significance level (P<0.05) is indicated with a *.

### SI 4: Results of the generalized linear mixed effects models measuring the association between gut microbiome diversity and survival to following breeding season

Summaries of the model effects for A) all collared flycatchers, B) male collared flycatchers, and C) female collared flycatchers and ANOVA type III Wald's chi-square test tables for D) all collared flycatchers, E) male collared flycatchers, and F) female collared flycatchers.

| **Predictor** | **estimate** | **s.e.** | **z** | **P** | | **estimate** | **s.e.** | **z** | **P** |
|---|---|---|---|---|---|---|---|---|---|
| **A** | | | | | | | | | |
| **Shannon** | 0.368 | 0.221 | 1.664 | 0.096 | **Chao1** | <0.001 | <0.001 | 0.435 | 0.664 |
| **Sex (male)** | 0.137 | 0.400 | 0.342 | 0.732 | | 0.013 | 0.388 | 0.033 | 0.973 |
| **Body condition** | -0.261 | 0.352 | -0.741 | 0.459 | | -0.253 | 0.350 | -0.721 | 0.471 |
| **Age group 1.5-4 yrs** | 0.494 | 0.531 | 0.930 | 0.352 | | 0.595 | 0.525 | 1.134 | 0.257 |
| **Age group 4+ yrs** | 0.063 | 0.879 | 0.072 | 0.943 | | 0.213 | 0.869 | 0.246 | 0.806 |
| **B** | | | | | | | | | |
| **Shannon** | 1.236 | 0.669 | 1.848 | 0.065 | **Chao1** | -0.001 | -0.003 | -0.266 | 0.790 |
| **Body condition** | 1.137 | 0.669 | 1.848 | 0.065 | | 0.588 | 0.675 | 0.871 | 0.384 |
| **Age group 1.5-4 yrs** | 17.137 | 836.093 | 0.020 | 0.984 | | 23.500 | 2048.00 | 0.011 | 0.991 |
| **Age group 4+ yrs** | 17.772 | 836.093 | 0.021 | 0.983 | | 23.830 | 2048.00 | 0.012 | 0.991 |
| **C** | | | | | | | | | |
| **Shannon** | -0.010 | 0.809 | -1.252 | 0.211 | **Chao1** | <0.001 | 0.106 | 0.745 | 0.457 |
| **Body condition** | -0.687 | 0.435 | -1.579 | 0.114 | | -0.642 | 0.433 | -1.482 | 0.138 |
| **Age group 1.5-4 yrs** | 0.011 | 0.588 | 0.019 | 0.985 | | 0.050 | 0.585 | 0.085 | 0.932 |
| **Age group 4+ yrs** | -33.700 | 1.357 | <0.000 | >0.999 | | -32.210 | 6.642 | <0.001 | >0.999 |
| | Chisq. | df | P | | | Chisq. | df | P | |
| **D** | | | | | | | | | |
| **Shannon** | 2.768 | 1 | 0.096 | | **Chao1** | 0.189 | 1 | 0.664 | |
| **Sex** | 0.117 | 1 | 0.732 | | | 0.001 | 1 | 0.973 | |
| **Body condition** | 0.549 | 1 | 0.459 | | | 0.520 | 1 | 0.471 | |
| **Age group** | 1.100 | 2 | 0.577 | | | 1.443 | 2 | 0.486 | |
| **E** | | | | | | | | | |
| **Shannon** | 3.416 | 1 | 0.065 | | **Chao1** | 0.071 | 1 | 0.790 | |
| **Body condition** | 1.983 | 1 | 0.159 | | | 0.758 | 1 | 0.394 | |
| **Age group** | 0.421 | 2 | 0.810 | | | 0.131 | 2 | 0.937 | |
| **F** | | | | | | | | | |
| **Shannon** | 0.985 | 1 | 0.321 | | **Chao1** | 0.554 | 1 | 0.457 | |
| **Body condition** | 2.495 | 1 | 0.114 | | | 2.198 | 1 | 0.138 | |
| **Age group** | <0.001 | 2 | 0.999 | | | 0.007 | 2 | 0.996 | |

* Significance level (P<0.05) is indicated with a *.

### SI 5: Results of the permutational multivariate analysis of variance measuring whether ARS, survival to following breeding season and LRS contribute to compositional differences in gut microbiome beta diversity.

Summary tables of the models for permutational multivariate analysis of variance (PERMANOVA) measuring whether A) ARS, B) survival to following breeding season, and C) LRS contribute to compositional differences in gut microbiome beta diversity. Other explanatory factors in each model include body condition, age group, area, and hatch date. In the models in which both sexes are included, sex is used as an explanatory factor.

A)

| **Predictor** | **df** | **SumOfSq.** | **R2** | **F** | **P** |
|---|---|---|---|---|---|
| **<u>BOTH SEXES</u>** | | | | | |
| **ARS** | 1 | 0.545 | 0.010 | 1.282 | 0.118 |
| **Sex** | 1 | 0.602 | 0.011 | 1.414 | 0.074 |
| **Body condition** | 1 | 0.358 | 0.007 | 0.841 | 0.760 |
| **Age group** | 2 | 0.763 | 0.014 | 0.897 | 0.707 |
| **Area** | 9 | 4.478 | 0.083 | 1.170 | ***0.025**** |
| **Hatch date** | 1 | 0.377 | 0.007 | 0.887 | 0.669 |
| **Residual** | 110 | 46.803 | 0.868 | | |
| **<u>MALES</u>** | | | | | |
| **ARS** | 1 | 0.562 | 0.031 | 1.400 | 0.089 |
| **Body condition** | 1 | 0.431 | 0.023 | 1.073 | 0.327 |
| **Age group** | 2 | 1.165 | 0.063 | 1.450 | ***0.028**** |
| **Area** | 4 | 1.859 | 0.101 | 1.157 | 0.108 |
| **Hatch date** | 1 | 0.351 | 0.019 | 0.874 | 0.608 |
| **Residual** | 35 | 14.056 | 0.763 | | |
| **<u>FEMALES</u>** | | | | | |
| **ARS** | 1 | 0.459 | 0.013 | 1.066 | 0.308 |

| | | | | | |
|---|---|---|---|---|---|
| **Body condition** | 1 | 0.324 | 0.009 | 0.752 | 0.923 |
| **Age group** | 2 | 0.717 | 0.021 | 0.833 | 0.901 |
| **Area** | 9 | 4.592 | 0.132 | 1.185 | ***0.012**** |
| **Hatch date** | 1 | 0.394 | 0.011 | 0.914 | 0.637 |
| **Residual** | 66 | 28.411 | 0.814 | | |

B)

| **Predictor** | **df** | **SumOfSq.** | **R2** | **F** | **P** |
|---|---|---|---|---|---|
| **BOTH SEXES** | | | | | |
| **Survival** | 1 | 0.499 | 0.008 | 1.168 | 0.190 |
| **Sex** | 1 | 0.657 | 0.011 | 1.536 | ***0.032**** |
| **Body condition** | 1 | 0.339 | 0.006 | 0.792 | 0.845 |
| **Age group** | 2 | 0.769 | 0.013 | 0.899 | 0.700 |
| **Area** | 10 | 4.860 | 0.081 | 1.137 | ***0.042**** |
| **Residual** | 123 | 52.599 | 0.881 | | |
| **MALES** | | | | | |
| **Survival** | 1 | 0.332 | 0.018 | 0.806 | 0.768 |
| **Body condition** | 1 | 0.420 | 0.022 | 1.020 | 0.379 |
| **Age group** | 2 | 1.157 | 0.061 | 1.406 | 0.051 |
| **Area** | 4 | 1.689 | 0.090 | 1.026 | 0.401 |
| **Residual** | 37 | 15.223 | 0.809 | | |
| **FEMALES** | | | | | |
| **Survival** | 1 | 0.493 | 0.012 | 1.142 | 0.206 |
| **Body condition** | 1 | 0.288 | 0.007 | 0.668 | 0.991 |
| **Age group** | 2 | 0.805 | 0.200 | 0.934 | 0.670 |
| **Area** | 10 | 5.028 | 0.125 | 1.166 | ***0.016**** |
| **Residual** | 78 | 33.632 | 0.836 | | |

C)

| **Predictor** | **df** | **SumOfSq.** | **R2** | **F** | **P** |
|---|---|---|---|---|---|
| **BOTH SEXES** | | | | | |
| **LRS** | 1 | 0.395 | 0.007 | 0.925 | 0.565 |
| **Sex** | 1 | 0.606 | 0.011 | 1.420 | 0.071 |
| **Body condition** | 1 | 0.379 | 0.007 | 0.887 | 0.650 |
| **Age group** | 2 | 0.807 | 0.150 | 0.945 | 0.580 |
| **Area** | 9 | 4.434 | 0.082 | 1.155 | ***0.022**** |
| **Hatch date** | 1 | 0.373 | 0.007 | 0.873 | 0.691 |
| **Residual** | 110 | 46.935 | 0.870 | | |
| **MALES** | | | | | |
| **LRS** | 1 | 0.450 | 0.024 | 1.102 | 0.281 |
| **Body condition** | 1 | 0.406 | 0.022 | 0.994 | 0.419 |
| **Age group** | 2 | 1.184 | 0.064 | 1.450 | ***0.038**** |
| **Area** | 4 | 1.752 | 0.095 | 1.073 | 0.285 |
| **Hatch date** | 1 | 0.348 | 0.019 | 0.852 | 0.653 |
| **Residual** | 35 | 18.423 | 0.775 | | |
| **FEMALES** | | | | | |
| **LRS** | 1 | 0.379 | 0.011 | 0.878 | 0.709 |
| **Body condition** | 1 | 0.321 | 0.009 | 0.743 | 0.948 |
| **Age group** | 2 | 0.740 | 0.021 | 0.858 | 0.849 |
| **Area** | 9 | 4.531 | 0.130 | 1.167 | ***0.016**** |
| **Hatch date** | 1 | 0.453 | 0.013 | 1.050 | 0.332 |
| **Residual** | 66 | 28.473 | 0.816 | | |

* Significance level (P<0.05) is indicated with a *.
